# Dopamine Release Neuroenergetics in Mouse Striatal Slices

**DOI:** 10.1101/2020.06.02.130328

**Authors:** Msema Msackyi, Yuanxin Chen, Wangchen Tsering, Ninghan Wang, Jingyu Zhao, Hui Zhang

## Abstract

Parkinson’s disease (PD) is the second most common neurodegenerative disease. Dopamine (DA) neurons in the substantia nigra par compacta with axonal projections to the dorsal striatum (dSTR) degenerate in PD while in contrast, DA neurons in the ventral tegmental area with axonal projections to the ventral striatum including the nucleus accumbens (NAcc) shell, are largely spared. To understand the pathogenesis of PD, it is important to study the neuroenergetics of DA neurons. This study aims to uncover the relative contribution of glycolysis and oxidative phosphorylation (OxPhos) to evoked DA release in the striatum. We measured evoked DA release in mouse striatal brain slices by fast-scan cyclic voltammetry every 2 minutes. Blocking OxPhos caused a greater reduction in evoked DA release in the dSTR compared to the NAcc shell, and blocking glycolysis caused a greater reduction in evoked DA release in the NAcc shell than in the dSTR. Furthermore, when glycolysis was bypassed in favor of direct OxPhos, evoked DA release in the NAcc shell was decreased by ∼50% over 40 minutes whereas evoked DA release in the dSTR was largely unaffected. These results demonstrated that the dSTR relies primarily on OxPhos for energy production to maintain evoked DA release whereas the NAcc shell relies more on glycolysis. Using two-photon imaging, we consistently found that the oxidation level of the DA terminals was higher in the dSTR than in the NAcc shell. Together, these findings partially explain the specific vulnerability of DA terminals in the dSTR to degeneration in PD.

**Significant statement:** The neuroenergetics of dopaminergic neuron is important to understand Parkinson’s disease (PD), a neurodegenerative disorder associated with mitochondrial dysfunctions. However, the relative contributions of glycolysis and oxidative phosphorylation (OxPhos) to presynaptic energy demands in DA terminals are unclear. We addressed this question by measuring DA release in the dorsal striatum and nucleus accumbens (NAcc) shell of mouse brain using FSCV under reagents blocking different energy systems. We found that the NAcc shell relies on both glycolysis and OxPhos to maintain DA release while the dSTR relies heavily on OxPhos. We demonstrate the different neuroenergetics of DA terminals in these two brain areas, providing new fundamentally important insight into the specific vulnerability of DA terminals in the dSTR to degeneration in PD.

## Introduction

Energy metabolism is crucial for the functioning of the brain, which requires energy in the form of ATP, to fuel the basic cellular functions, including maintaining ion gradients and cycling presynaptic synaptic vesicles (Ames, 2000). The major source of energy in the brain is glucose, which is broken down through glycolysis and oxidative phosphorylation (OxPhos) to produce ATP. Glycolysis, which occurs in the absence of oxygen, rapidly produces a net total of two ATP and two molecules of pyruvate from glucose. OxPhos, which occurs in mitochondria, is slower than glycolysis but has much higher yield of ATP production. OxPhos produces 32 ATP, and contributes roughly 80% of energy in the brain (Hall et al., 2012). The brain uses the majority of energy at the synapses and nodes of Ranvier, where mitochondria is abundant and the demand for energy is high.

The utilization of energy at the presynaptic compartments and energetic cost of presynaptic function has been of enormous interest (Ashrafi and Ryan, 2017). Recent studies in the mouse calyx of Held (Lujan et al., 2016) and in rat hippocampal cell culture (Sobieski et al., 2017) have attempted to parse the contribution of each of the energy production pathways, OxPhos and glycolysis, in glutamatergic neurons. Research conducted in mouse calyx of Held concluded that glycolysis has a greater effect on basal glutamatergic transmission than OxPhos (Lujan et al., 2016). Research conducted in single neuron rat hippocampal culture expanded on the findings of the previous study (Sobieski et al., 2017). The authors concluded that either OxPhos or glycolysis is sufficient for the glutamate transmission, however OxPhos was of primary importance for high demand transmission. In the absence of glycolysis, monocarboxylates (e.g., lactate) acts as alternative fuel source for oxidative phosphorylation. As anaerobic glycolysis occurs in astrocytes, lactate builds up and is subsequently shuttled to neurons through monocarboxylate transport (MCT) and used for fuel (Machler et al., 2016; Magistretti and Allaman, 2018). Although energy production in the brain is well understood, the dependence of DA release on the different energy production systems remains unclear.

DA neurons are tonically active with extensive arborization (Matsuda et al., 2009), and they are thought to be more metabolically expensive (Pissadaki and Bolam, 2013). This high metabolic demand could result in higher rates of energy production and oxidative stress. Cell culture research has shown that, compared to the VTA neurons, SNc DA neurons have a higher basal rate of mitochondrial OxPhos, a lower reserve capacity of spare ATP, a higher density of axonal mitochondria and oxidative stress, and a considerably more complex axonal arborization (Pacelli et al., 2015). However, to date, no one has studied the energetic cost of presynaptic DA neurotransmission. Studying the neuroenergetics of DA neurotransmission is particularly important in the context of neurodegenerative disorders, such as Parkinson’s disease (PD), which is characterized by mitochondrial dysfunction and oxidative stress (Surmeier et al., 2011; Bose and Beal, 2016). In PD, the DA terminals in the dSTR projected from SNc DA neurons degenerates whereas the DA terminals in the ventral striatum, including NAcc shell projected from VTA DA neurons, are largely spared. In this study, we investigated the role of glycolysis and OxPhos in energy production for evoked release of DA in the dSTR and the NAcc shell. We found that the dSTR relies more heavily on OxPhos for evoked DA release compared to the NAcc shell while, in contrast, the NAcc shell relies more on glycolysis for evoked DA release compared to the dSTR.

## Materials and Methods

### Animals and slice preparation

The use of the animals followed the National Institutes of Health guidelines and was approved by the Institutional Animal Care and Use Committee at Thomas Jefferson University. All efforts were made to minimize the number of animals used. Adult wild-type (WT) mice at the age of 2 to 10-months of both sexes were used. TH-mito-roGFP mice expressing a redox-sensitive variant of green fluorescent protein (roGFP) with a mitochondrial-matrix targeting sequence under the control of the TH promoter were obtained from Dr. Surmeier at Northwestern University (Guzman et al., 2010).

Mice were euthanized via cervical dislocation with no anesthetic. The brain was immediately removed from the skull. Coronal striatal brain slices at 300 μm were prepared on a vibratome (VT1200, Leica, Solms, Germany) for electrophysiological recording. The striatal slices were allowed to recover for at least 1 hr at room temperature (RT) in a holding chamber containing oxygenated artificial cerebrospinal fluid (ASCF) and then placed in a recording chamber superfused (1.5 mL/min) with ACSF (in mM: 125 NaCl, 2.5 KCl, 26 NaHCO3, 2.4 CaCl2, 1.3 MgSO4, 0.3 KH2PO4, and 10 glucose) at 36 °C. At least 3 mice were used per condition unless specified.

### Fast-scan cyclic voltammetry and amperometry

Fast-scan cyclic voltammetry (FSCV) was used to measure evoked DA release in the striatum. Electrochemical recording and electrical stimulations were performed as previously described (Zhang and Sulzer, 2003). Briefly, disk carbon fiber electrodes of 5 μm diameter with a freshly cut surface were placed into the dSTR or the NAcc shell ∼50 μm below the exposed surface. For FSCV, a triangular voltage wave (−450 to +800 mV at 280 V/sec vs Ag/AgCl) was applied to the recording electrode every 100 ms with a pulse duration of 8.5 ms and a ramp of 294 mV/ms via an Axopatch 200B (Axon Instruments, Foster City, CA). The current was recorded with an Axopatch 200B amplifier with a low-pass Bessel filter setting at 10 kHz, digitized at 25 kHz (ITC-18 board; InstruTech, Great Neck, NY). Triangular wave generation and data acquisition were controlled by a personal computer running a locally written (Dr. E. Mosharov, Columbia University, New York, NY) IGOR program (WaveMetrics, Lake Oswego, OR). Striatal slices were electrically stimulated by an Iso-Flex stimulus isolator triggered by a Master-8 pulse generator (A.M.P.I., Jerusalem, Israel) using a bipolar stimulating electrode placed at a distance of ∼100 μm from the recording electrode. Background-subtracted cyclic voltammograms served for electrode calibration and to identify the released substance.

### 2PLSM: mitochondrial roEGFP imaging in living brain

Striatal slices from TH-mito-roGFP transgenic mice were imaged at physiological temperatures (36 ° C). Optical imaging of roGFP signals were acquired using an Ultima multiphoton laser scanner (Bruker) for BX51/61 Olympus microscope with a Titanium-sapphire laser (Chameleon-Ultra2, Coherent Laser Group), equipped with a 20 × 1.0 NA water immersion objective (XLUMPLFL20XW, Olympus). Dual excitation beam (800nm/900nm) was used to excite roGFP protein respectively and roGFP fluorescence was collected at 490–560 nm. The ratio of 800 nm/900 nm was taken as an index of the oxidation level. Images were captured in 8-bit, 45 × 45 µm regions of interest at 512 × 512 pixel.

### Experimental Design

The slices were stimulated every 2 min in the dSTR or NAcc shell for 40 min and DA release was measured with FSCV. Pharmacological reagents that selectively inhibit glycolysis or OxPhos were added by perfusion on mouse brain slices. Experiments were generally initiated after three stimuli to confirm a consistent and robust response. Normalization was done in comparison to the average of the first three stimuli unless otherwise specified.

Energy production systems under various conditions are shown in the schematic and table (Fig. 1). Notably, although OxPhos produces roughly 16 times more ATP than glycolysis produces per glucose molecule, the process is much more time consuming than glycolysis. OxPhos has many more steps than glycolysis including two whole turns of the eight-step Krebs cycle per glucose molecule followed by ATP production via the mitochondria complex and the electron transport chain (Fig. 1A). The MCT system itself does not produce energy; it transports mainly lactate as fuel for OxPhos (Fig. 1A). The pathway leading from import of glucose or the breakdown of glycogen to pyruvate and then to lactate to be exported is the pathway astrocytes use for MCT (Magistretti and Allaman, 2018).

**Figure 1.**
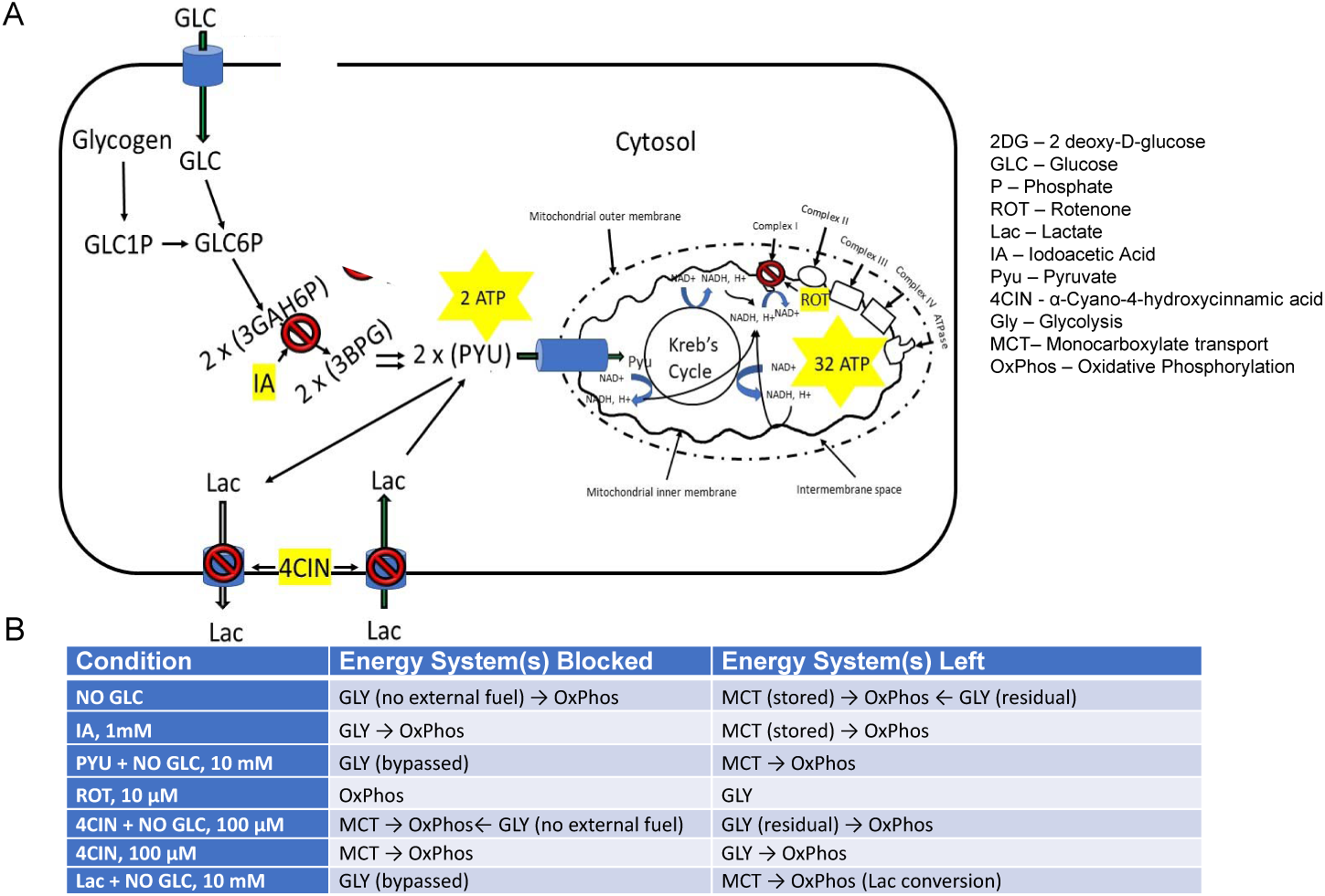
Schematics and table showing cellular energy production systems under various conditions. **A**, A schematic showing the different energy systems in the striatum with inhibitors to block the different energy production pathways. **B**, A table showing how the different conditions affect the various energy production systems related to evoked DA release.

### Statistical Analysis

Experimental values in the text and in the figures are mean ± SEM. The Student’s t-test was used for paired data and two-way ANOVA followed by Bonferroni post-test was performed using Prism 7.0 (GraphPad) to determine statistical significance for all grouped data unless otherwise specified. The difference was considered significant at levels of * < 0.05; ** < 0.01; *** p < 0.001; **** < 0.0001; n.s. stands for no significance, n stands for the number of experiments.

## Results

### Evoked DA release is higher in the dSTR than in the NAcc shell

To investigate evoked DA release in the dSTR and the NAcc shell, we measured DA release via a single-pulse electrical stimulus every 2 min for 40 min under regular glucose-containing ACSF using FSCV (Fig. 2A). Evoked DA release in the dSTR was significantly higher than that in the NAcc shell (Fig. 2B, C) with a dSTR:NAcc shell ratio of DA release of 1:0.71 (Fig. 2C). Evoked DA release in both the dSTR and the NAcc shell is stable over 40 min showing no significant difference with an insignificant 5% run-down (Fig. 2D).

**Figure 2.**
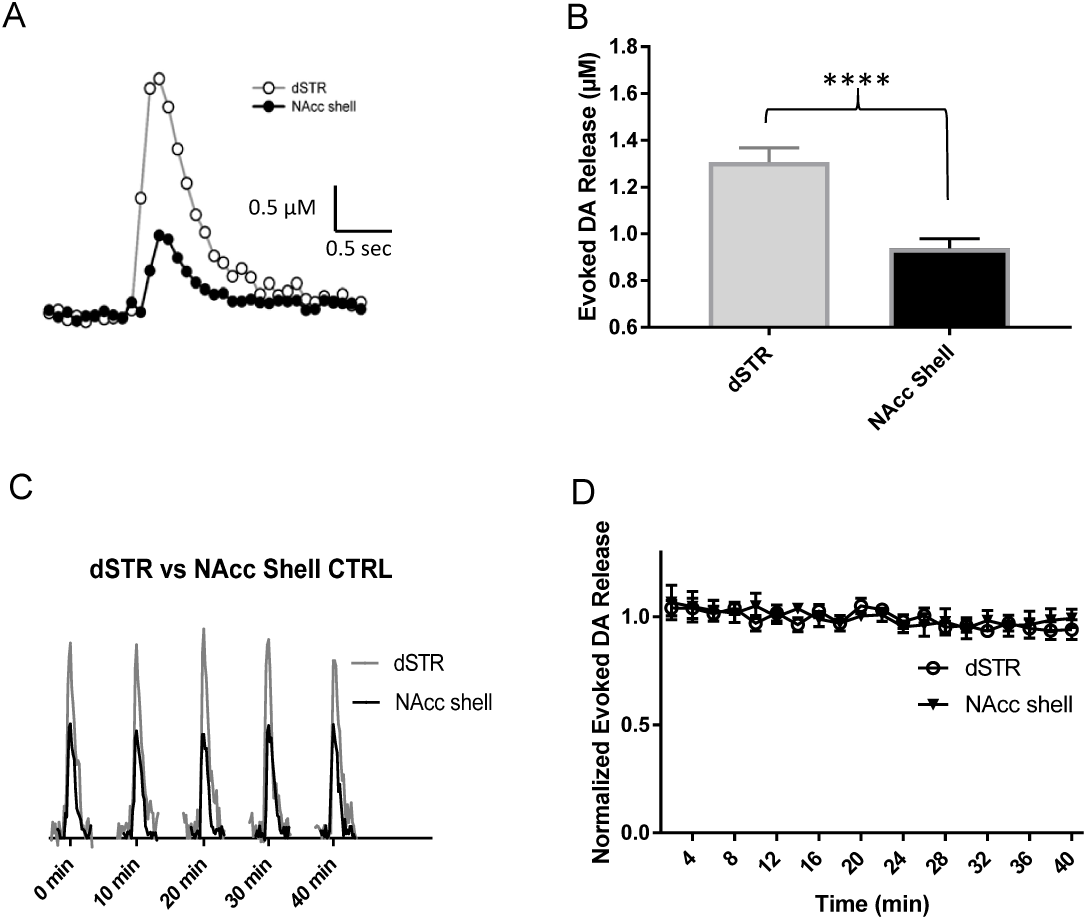
Evoked DA release is higher in the dSTR than in the NAcc shell. Evoked DA release in the dSTR is significantly higher than that in the NAcc shell. **A**, Representative FSCV traces of 1p-stimulation-evoked DA release recorded in the dSTR and the NAcc shell. **B**, Plot showing average amplitudes of 1p-evoked DA release (dSTR, n = 8; NAcc shell, n = 9 NAcc shell, *p* < *0.0001*, Student t-test). **C**, Representative traces of 1p-evoked DA release over 40 min taken every 10 min under control conditions. **D**, Normalized evoked DA release over 40 min measured every 2 min in the dSTR and the NAcc shell (dSTR, n = 8; NAcc shell, n = 9, n.s., two-way ANOVA with Bonferroni *post hoc* test).

### Similar diminishment of evoked DA release in the dSTR and the NAcc shell under glucose-deprived conditions

To test the evoked DA release in the dSTR relative to the NAcc shell under conditions of energy-deprivation, we perfused striatal slices with ACFS containing no glucose. Under conditions of glucose-deprivation, the evoked DA release in both areas decreased over time with a significant diminishment compared to their respective controls (Fig. 3A,B). The diminishment of the evoked DA release in the dSTR was similar to the NAcc shell (Fig. 3C). Under these conditions, no external fuel was perfused in the slices that could be used for energy production.

**Figure 3.**
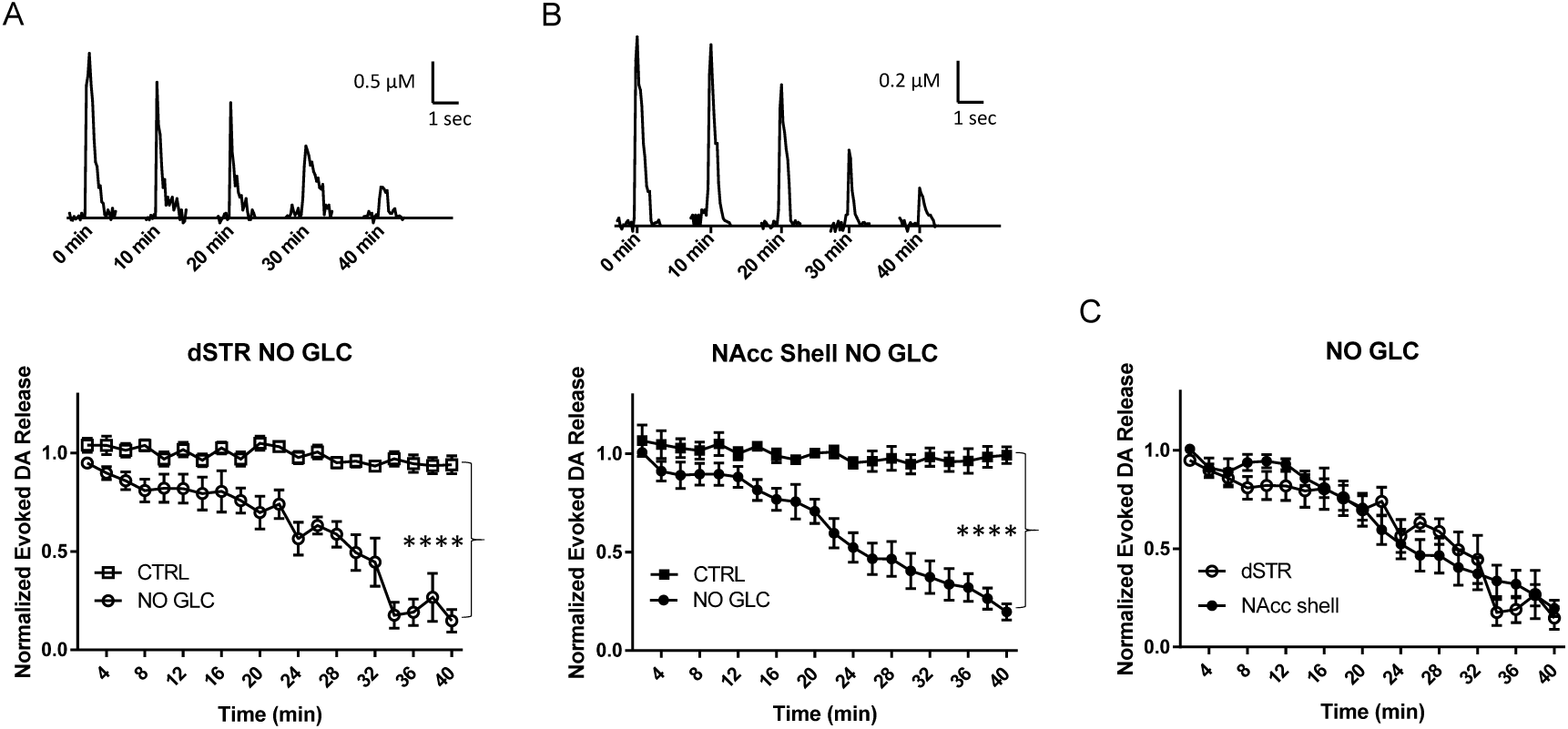
Similar diminishment of evoked DA release in the dSTR and the NAcc shell under glucose-deprived conditions. **A**, Representative traces of 1p-evoked DA release in the dSTR over 40 min taken every 10 min under glucose-deprived conditions (NO GLC, top) and plot of the normalized evoked DA release over 40 min measured every 2 min under glucose-deprived conditions (bottom, NO GLC, n = 11; control (CTRL), n = 8, NO GLC *vs* CTRL, *p* < *0.0001*, two-way ANOVA with Bonferroni *post hoc* test). **B**, Representative traces of 1p-evoked DA release in the NAcc shell over 40 min taken every 10 min under glucose-deprived conditions (top) and plot of the normalized evoked DA release over 40 min measured every 2 min under glucose-deprived conditions (bottom, NO GLC, n = 8; CTRL, n = 9, NO GLC *vs* CTRL, *p* < *0.0001*, two-way ANOVA with Bonferroni *post hoc* test). **C**, Comparison of normalized evoked DA release in the dSTR and NAcc shell over 40 min under glucose-deprived conditions showing no statistical significance, dSTR *vs* NAcc shell, n.s., two-way ANOVA with Bonferroni *post hoc* test.

### Glycolysis inhibition diminishes evoked DA release in the NAcc shell more rapidly than in the dSTR

We then perfused the brain slices with glycolysis inhibitor iodoacetate (IA, 1 mM) to investigate the importance of glycolytic energy production for evoked DA release in the striatum. IA blocks glycolysis by hindering the conversion of GAH6P to BPG (Fig. 1A). IA was used under regular ACFS conditions. Under these conditions, evoked DA in the dSTR and the NAcc shell diminished by approximately 90% over 40 min (Fig. 4A, B). The diminishment in evoked DA release occurred more rapidly in the NAcc shell when compared to the dSTR (Fig. 4C). This indicates that while glucose is present, the NAcc shell relies more heavily on glucose than the dSTR does for evoked DA release. Evoked DA release in the dSTR was similar in this condition compared to glucose-deprived conditions over 40 min perfusion (Fig. 4D), however, evoked DA release in the NAcc shell, was significantly lower in this condition than under glucose-deprived conditions (Fig. 4E). This suggests that presence of glucose is associated with an increased dependence on glycolysis for evoked DA release in the NAcc shell. In contrast, evoked DA release in the dSTR is similarly dependent on glycolysis regardless of whether glucose is present or not. Thus, the dSTR relies more on non-glycolytic energy than the NAcc shell when glucose is present.

**Figure 4.**
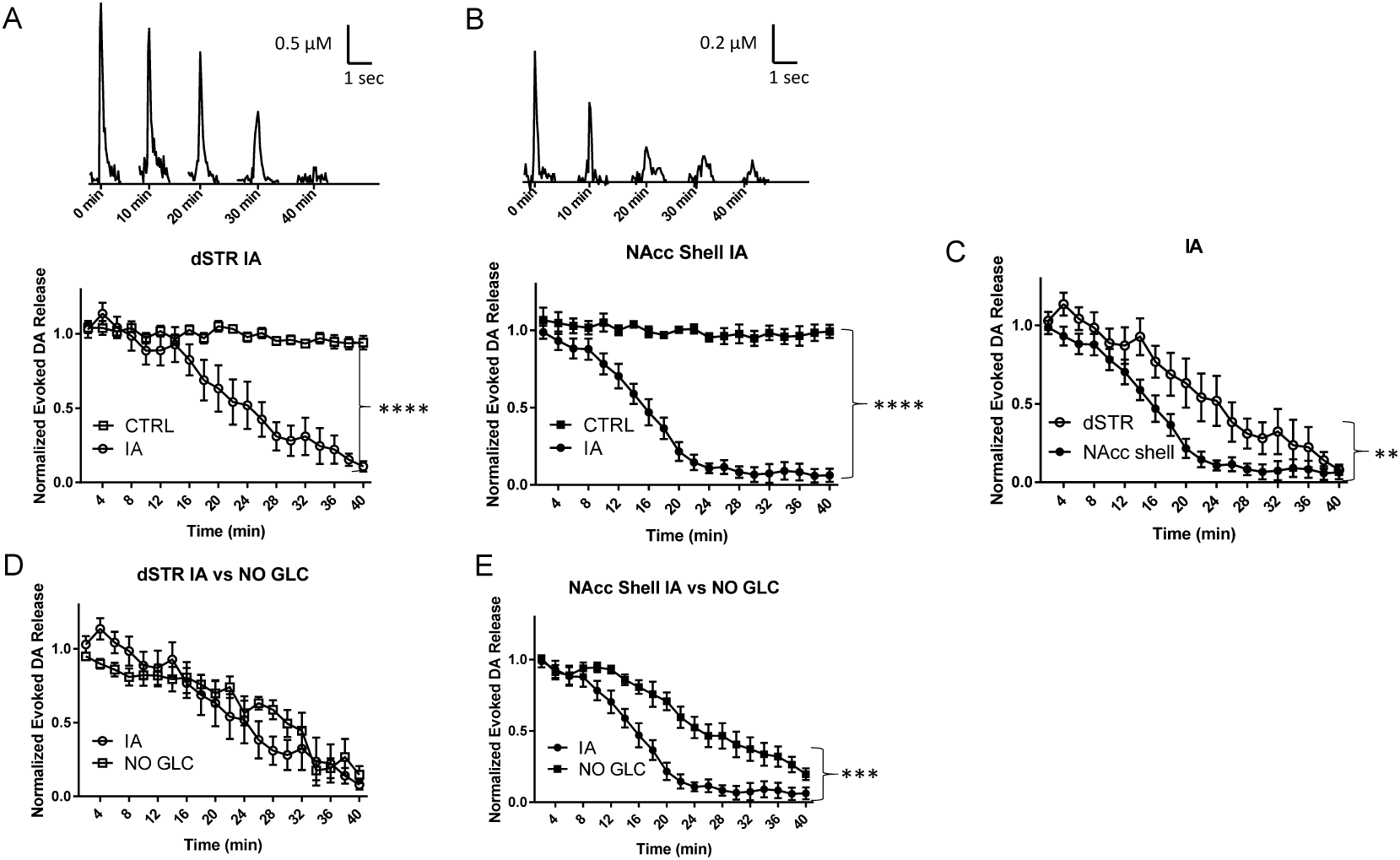
Glycolysis inhibition diminishes evoked DA release in the NAcc shell more rapidly than in the dSTR. **A**, Representative traces of 1p-evoked DA release in the dSTR over 40 min taken every 10 min when glycolysis was inhibited with 1 mM IA (top) and plot of the normalized evoked DA release over 40 min measured every 2 min (bottom, IA, n = 8; CTRL, n = 8, IA *vs* CTRL, *p* < *0.0001*, two-way ANOVA with Bonferroni *post hoc* test). **B**, Representative traces of 1p-evoked DA release in the NAcc over 40 min taken every 10 min when glycolysis was inhibited with 1 mM IA (top) and plot of the normalized evoked DA release over 40 min measured every 2 min (bottom, IA, n = 7; CTRL, n = 9, IA *vs* CTRL, *p* < *0.0001*, two-way ANOVA with Bonferroni *post hoc* test). **C**, Comparison of normalized evoked DA release in the dSTR and NAcc shell over 40 min under glycolysis inhibition conditions, dSTR *vs* NAcc shell, *p* < *0.01*, two-way ANOVA with Bonferroni *post hoc* test. **D**, Comparison of normalized evoked DA release in the dSTR over 40 min under glycolysis inhibition and glucose-deprived conditions showing no significant difference, IA, n = 8, NO GLC, n = 11, IA *vs* NO GLC, *p* < *0.01*, two-way ANOVA with Bonferroni *post hoc* test. **E**, Comparison of normalized evoked DA release in the NAcc over 40 min under glycolysis inhibition and glucose-deprived conditions showing glycolysis inhibition diminishes evoked DA release more rapidly than glucose-deprived condition, IA, n = 7; NO GLC, n = 8, IA *vs* NO GLC, *p* < *0.001*, two-way ANOVA with Bonferroni *post hoc* test.

### Bypassing glycolysis diminishes evoked DA release in the NAcc shell more rapidly than the dSTR

At the end of glycolysis, glucose is converted into two molecules of pyruvate which feed the Krebs cycle and lead to OxPhos (Fig. 1A). Pyruvate is therefore important for OxPhos and can bypass glycolysis to produce ATP. This is important in determining how well OxPhos alone can maintain evoked DA release. Under glucose-deprived conditions, 10 mM pyruvate was perfused and evoked DA release in the dSTR was measured. Evoked DA release was slightly diminished for the first 6 min, and then it rebounded, and by the end of the perfusion, evoked DA release was the same compared to the control condition (Fig. 5A). This initial diminishment could be caused by glycolysis being bypassed, hence the quick production of ATP from glycolysis would be abrogated. This dynamism did not occur for evoked DA release in the NAcc shell. The evoked release in the NAcc shell kept decreasing and was diminished by roughly 50% over 40 min (Fig. 5B). Aside from an initial diminishment, the evoked DA release in the dSTR was maintained at control levels whereas the evoked DA release in the NAcc shell dropped to half of the control levels (Fig. 5C). This indicates evoked DA release in the dSTR is more reliant on OxPhos than the NAcc shell.

**Figure 5.**
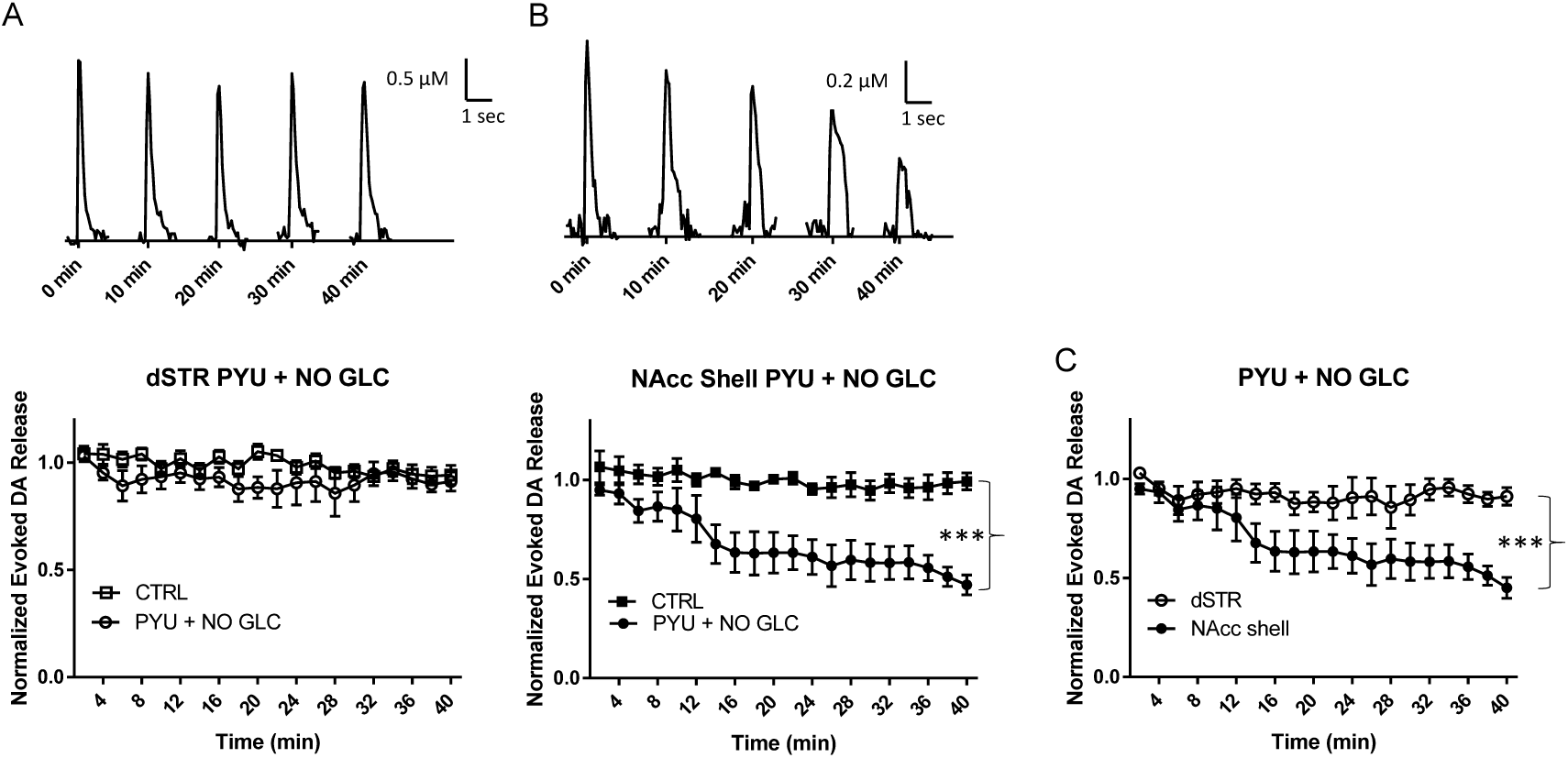
Bypassing glycolysis diminishes evoked release of DA in the NAcc shell more than the dSTR. **A**, Representative traces of 1p-evoked DA release in the dSTR over 40 min taken every 10 min under when 10 mM pyruvate was administered under glucose-deprived conditions (PYU + NO GLC, top) and plot of the normalized evoked DA release over 40 min measured every 2 min (bottom, n = 9, CTRL; n = 8, PYU + NO GLC *vs* CTRL, *n.s.*, two-way ANOVA with Bonferroni *post hoc* test). **B**, Representative traces of 1p-evoked DA release in the NAcc shell over 40 min taken every 10 min under glucose-deprived conditions (top) and plot of the normalized evoked DA release over 40 min measured every 2 min (bottom, n = 7; CTRL, n = 9, PYU + NO GLC *vs* CTRL, *p* < *0.001*, two-way ANOVA with Bonferroni *post hoc* test). **C**, Comparison of normalized evoked DA release in the dSTR and NAcc shell over 40 min under condition bypassing glycolysis, dSTR *vs* NAcc shell, *p* < *0.001*, two-way ANOVA with Bonferroni *post hoc* test.

### Specific inhibition of OxPhos diminishes evoked DA release in the dSTR more than the NAcc shell

We then assessed the effect of direct inhibiting OxPhos on the evoked DA release in these brain areas by using rotenone (ROT), an inhibitor of complex I of the mitochondrial electron transport chain (Fig 1A). When 10 μM ROT was applied to the brain slices, evoked DA release in both the dSTR and the NAcc shell was significantly diminished (Fig. 6A,B), but the diminishment in the dSTR was quicker and significantly greater than that in the NAcc shell (∼55% vs. ∼30%, p < 0.001, Fig. 6C). Together with the condition of bypassing glycolysis using pyruvate (Fig. 5C), these results suggest that the dSTR is more reliant than NAcc on OxPhos. Furthermore, these results, in combination with the results from Fig. 4C, indicate that the NAcc is more reliant than dSTR on glycolysis for DA release.

**Figure 6.**
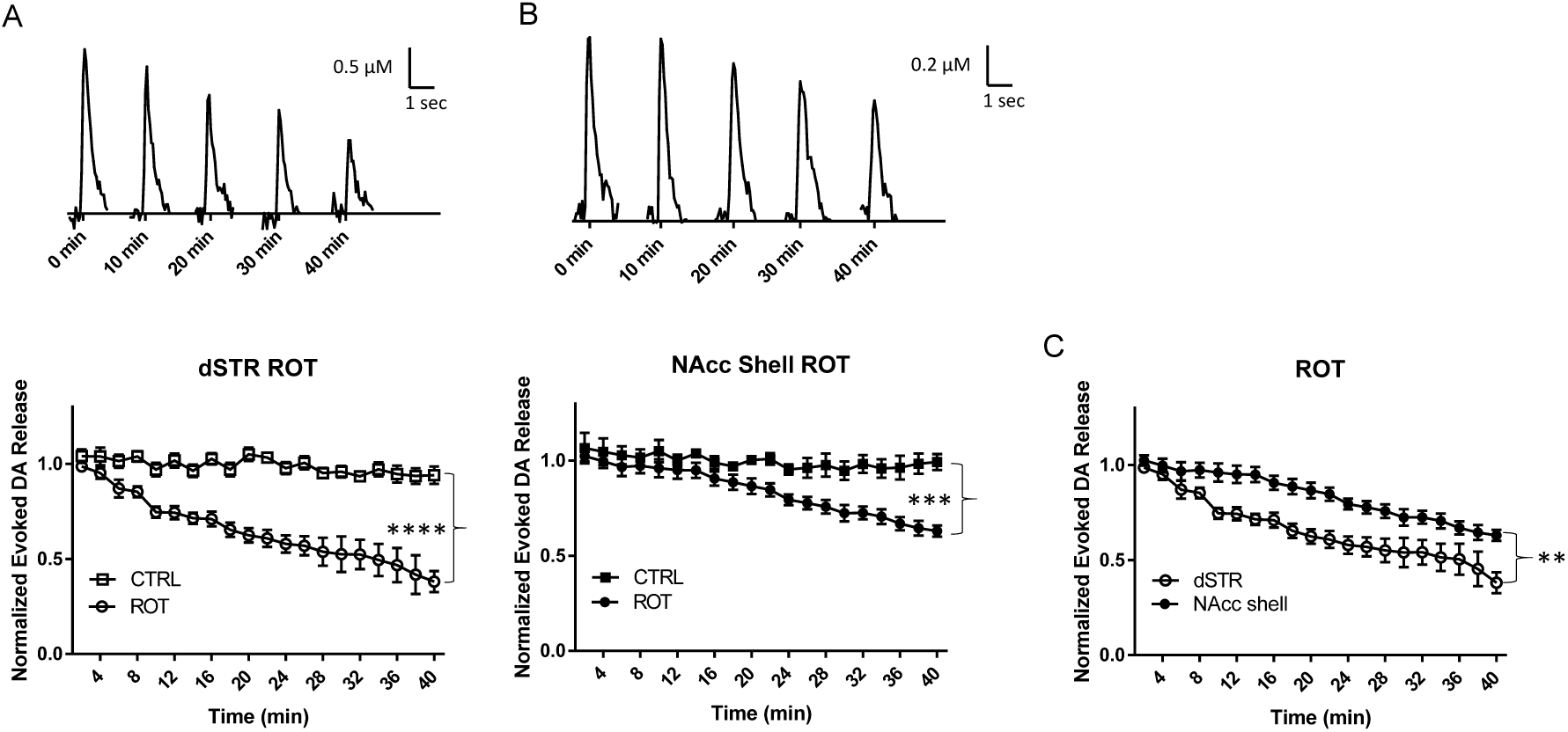
Specific inhibition of OxPhos diminishes evoked release of DA in the dSTR more than the NAcc shell. **A**, Representative traces of 1p-evoked DA release in the dSTR over 40 min taken every 10 min when complex I inhibited with 10 µM rotenone (ROT, top) and plot of the normalized evoked DA release over 40 min measured every 2 min (bottom, n = 6; CTRL, n = 8, ROT *vs* CTRL, *p* < *0.0001*, two-way ANOVA with Bonferroni *post hoc* test). **B**, Representative traces of 1p-evoked DA release in the NAcc shell over 40 min taken every 10 min when complex I inhibited with 10 µM rotenone (ROT, top) and plot of the normalized evoked DA release over 40 min measured every 2 min (bottom, n = 9; CTRL, n = 9, ROT *vs* CTRL, *p* < *0.001*, two-way ANOVA with Bonferroni *post hoc* test). **C**, Comparison of normalized evoked DA release in the dSTR and NAcc shell over 40 min under specific inhibition of OxPhos, dSTR *vs* NAcc shell, *p* < *0.01*, two-way ANOVA with Bonferroni *post hoc* test.

### MCT inhibition reduces evoked DA release in the dSTR but not in the NAcc shell

The MCT system may play a significant role in energy production especially under conditions of high energy demands which increase glucose consumption, mostly in astrocytes (Wyss et al., 2011; Nortley and Attwell, 2017; Brooks, 2018; Magistretti and Allaman, 2018). Astrocytes mainly fuel neurons via glucose metabolites, such as lactate or pyruvate (Lac: PYU = 10:1), which are then shuttled through MCTs to neighboring neurons (Machler et al., 2016; Magistretti and Allaman, 2018). To examine the role of MCT system and their relative importance to energy production in DA terminals, slices were perfused with 100 μM α-cyano-4-hydroxycinnamic acid (4CIN), a non-specific inhibitor of the MCT system (Fig. 1A). When 4CIN was administered under regular glucose conditions, evoked DA release in the dSTR was slightly but significantly diminished as compared to the control (Fig. 7A). This suggests that under normal conditions dSTR utilizes the MCT system as a fuel for energy production but not at a high level. In contrast, there was no significant diminishment of evoked DA release in the NAcc shell (Fig. 7B). There was an initial decrease in evoked DA release in the NAcc shell, but the diminishment was recovered over 40 min of perfusion. Interestingly, evoked DA release in the nAcc shell became highly variable during the course. Since glycolytic cells produce a high level of lactate that is exported under normal conditions, this variability in evoked DA release could be due to a buildup of lactate waste since the export of lactate was blocked, leading to acidification. In summary, the MCT system fuels evoked DA release in the dSTR slightly but not in the NAcc shell when glucose is present (Fig. 7C).

**Figure 7.**
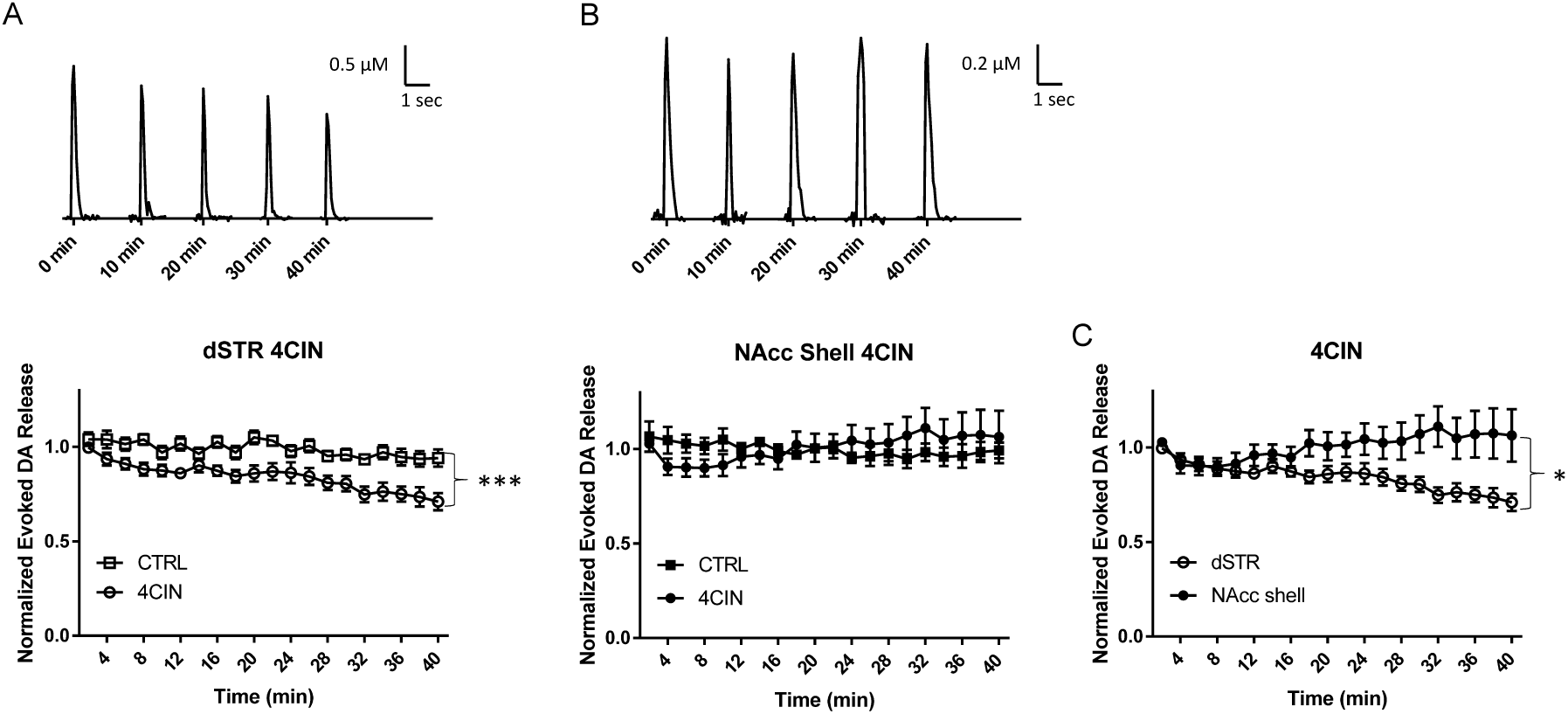
MCT inhibition reduces evoked DA release in the dSTR but not in the NAcc shell. **A**, Representative traces of 1p-evoked DA release in the dSTR over 40 min taken every 10 min when MCT inhibited with 100 µM 4CIN (top) and plot of the normalized evoked DA release over 40 min measured every 2 min (bottom, n = 10; CTRL, n = 8, 4CIN *vs* CTRL, *p* < *0.001*, two-way ANOVA with Bonferroni *post hoc* test). **B**, Representative traces of 1p-evoked DA release in the NAcc shell over 40 min taken every 10 min MCT inhibited with 100 µM 4CIN (top) and plot of the normalized evoked DA release over 40 min measured every 2 min (bottom, n = 9, CTRL, n = 9, 4CIN *vs* CTRL, *n.s*, two-way ANOVA with Bonferroni *post hoc* test). **C**, Comparison of normalized evoked DA release in the dSTR and NAcc shell over 40 min under inhibition of MCT, dSTR *vs* NAcc shell, *p* < *0.05*, two-way ANOVA with Bonferroni *post hoc* test.

### MCT inhibition under glucose-deprived conditions reduces evoked DA release in the striatum more rapidly compared to glucose-deprived conditions

To investigate the MCT system under energy-deprived conditions, slices were perfused with 100 μM 4CIN under glucose-deprived conditions. There is evidence that energy deprivation increases MCT from astrocytes (Schurr et al., 1988; Wyss et al., 2011; Magistretti and Allaman, 2018). Under glucose-deprived conditions, evoked DA release diminished more rapidly compared to glucose-deprivation alone in both region (Fig. 8A, B). The diminishment of evoked DA release in the dSTR and the NAcc shell was similar in response to 4CIN with no significant difference in these two areas (Fig. 8C). This suggests that both the dSTR and the NAcc shell rely on energy produced by the MCT system in the absence of glucose.

**Figure 8.**
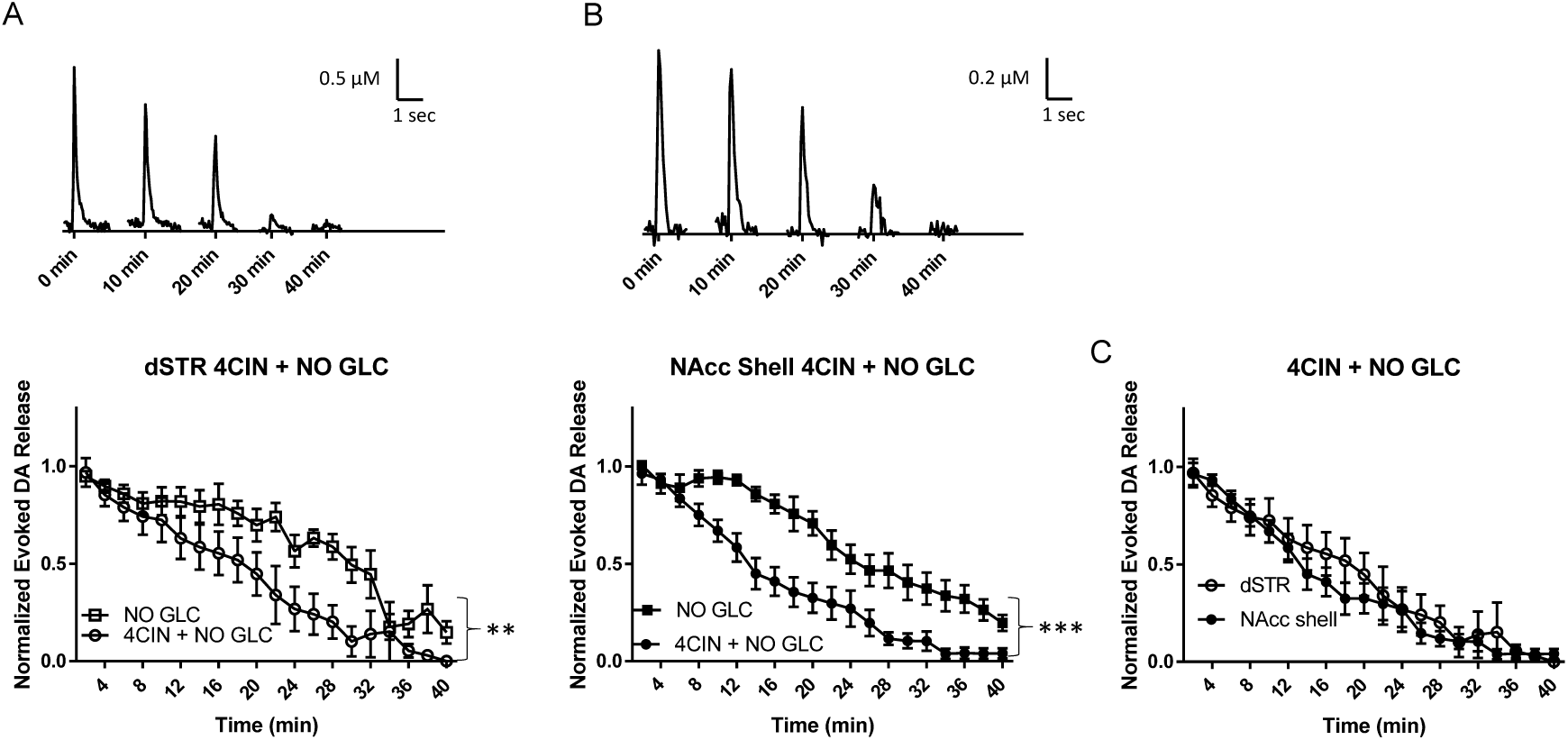
MCT inhibition under glucose-deprived conditions reduces evoked DA release in the striatum more rapidly compared to glucose-deprived conditions. Representative traces of 1p-evoked DA release in the dSTR over 40 min taken every 10 min when MCT inhibited with 100 µM 4CIN under glucose-deprived conditions (top) and plot of the normalized evoked DA release over 40 min measured every 2 min (bottom, n = 9; NO GLC, n = 11, 4CIN + NO GLC *vs.* NO GLC, *p* < *0.01*, two-way ANOVA with Bonferroni *post hoc* test). **B**, Representative traces of 1p-evoked DA release in the dSTR over 40 min taken every 10 min when MCT inhibited with 100 µM 4CIN under glucose-deprived conditions (top) and plot of the normalized evoked DA release over 40 min measured every 2 min (bottom, n = 9; NO GLC, n = 8, 4CIN + NO GLC *vs* NO GLC, *p* < *0.001*, two-way ANOVA with Bonferroni *post hoc* test). **C**, Comparison of normalized evoked DA release in the dSTR and NAcc shell over 40 min under MCT inhibition under glucose-deprived conditions, dSTR *vs* NAcc shell, *n.s*, two-way ANOVA with Bonferroni *post hoc* test.

### Energy production via the MCT system maintains evoked DA release at a higher level in the dSTR than in the NAcc shell

Considering that lactate is the main glucose metabolite that is shuttled through the MCT system to neurons (Machler et al., 2016; Magistretti and Allaman, 2018), slices were perfused with 10 mM lactate under glucose-deprived conditions. This can serve as another measurement of OxPhos energy production fueled by MCT system. This condition is complementary to measure the effect of inhibiting MCT on the evoked DA release (Fig. 8). Lactate can be used as fuel for OxPhos after it is converted to pyruvate (Fig. 1A). Lactate, when perfused under glucose-deprived conditions, led to an initial suppression of evoked DA release in the dSTR (Fig. 9A), but this diminishment largely rebounded over 40 min of perfusion. In contrast, the diminishment was more drastic in the NAcc shell and did not rebound (Fig. 9B). Evoked DA release was diminished to a greater degree in the NAcc shell when compared to the dSTR (Fig. 9C). Lactate and pyruvate conditions were similar but the initial diminishment in evoked DA release in the dSTR was longer with lactate, possibly due to the conversion of lactate to pyruvate which then fuels OxPhos.

**Figure 9.**
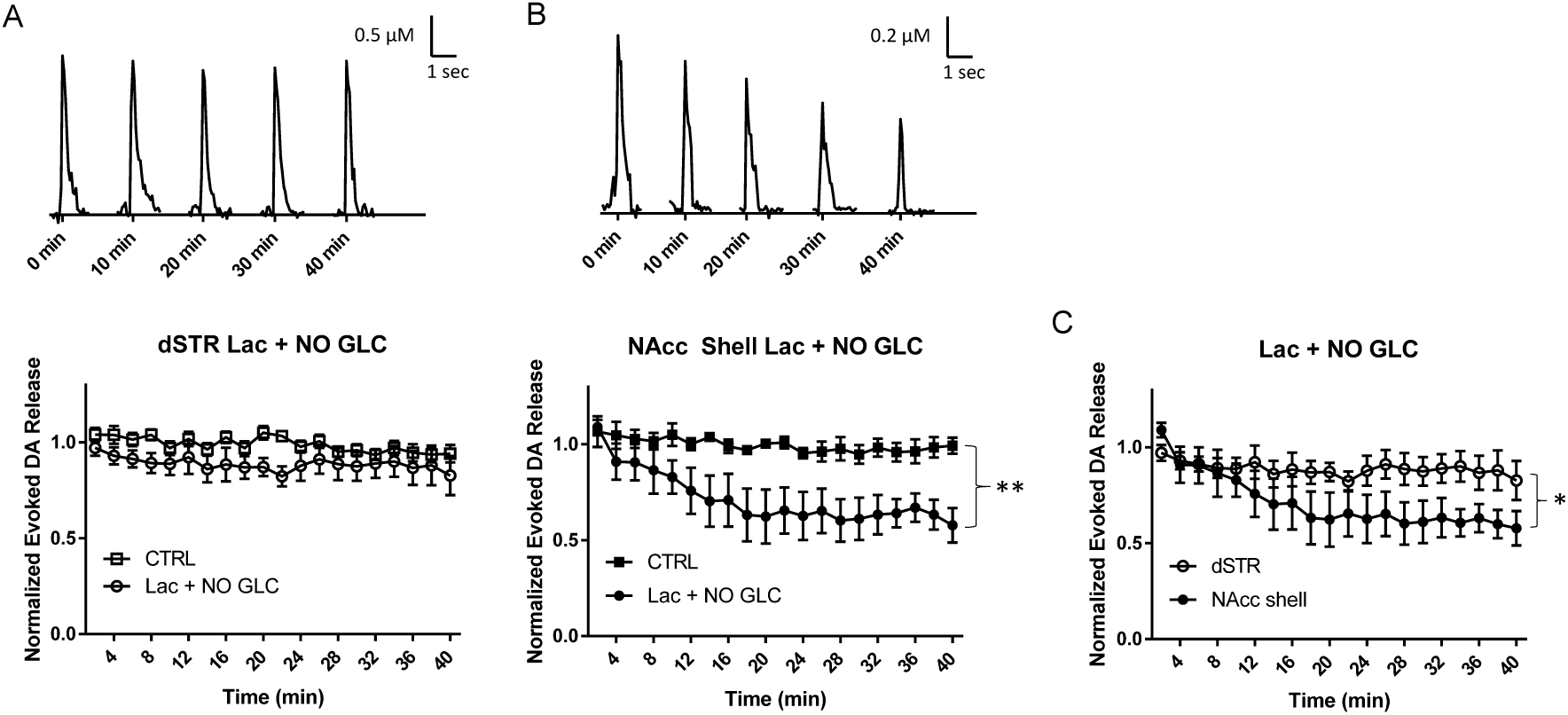
Energy production via the MCT system can maintain evoked DA release in the dSTR but not in the NAcc shell. Representative traces of 1p-evoked DA release in the dSTR over 40 min taken every 10 min when 10 mM lactate was administered under glucose-deprived condition (Lac + NO GLC, top) and plot of the normalized evoked DA release over 40 min measured every 2 min (bottom, n = 8; CTRL, n = 8, Lac + NO GLC *vs* CTRL, *n.s.*, two-way ANOVA with Bonferroni *post hoc* test). **B**, Representative traces of 1p-evoked DA release in the NAcc shell over 40 min taken every 10 min when 10 mM lactate was administered under glucose-deprived condition (Lac + NO GLC, top) and plot of the normalized evoked DA release over 40 min measured every 2 min (bottom, n = 7; CTRL, n = 9, Lac + NO GLC *vs* CTRL, *p* < *0.01*, two-way ANOVA with Bonferroni *post hoc* test). **C**, Comparison of normalized evoked DA release in the dSTR and NAcc shell over 40 min when 10 mM lactate was administered under glucose-deprived condition, dSTR *vs* NAcc shell, *p* < *0.05*, two-way ANOVA with Bonferroni *post hoc* test.

### Higher oxidation level in the DA terminals in the dSTR than that in the NAcc

The results present here show that in general evoked DA release in the dSTR largely depends on OxPhos energy production while evoked DA release in the NAcc shell more depends on glycolytic energy production. The byproducts of OxPhos are reactive species such as reactive oxygen species that can lead to oxidative stress (Bose and Beal, 2016). To directly assess whether there is a higher oxidation level in the dSTR, we measured the oxidation level using two-photon imaging of TH-mito-roGFP mice (Guzman et al., 2010). As expected, the oxidation level was higher in the dSTR than that in the NAcc (Fig. 10).

**Figure 10.**
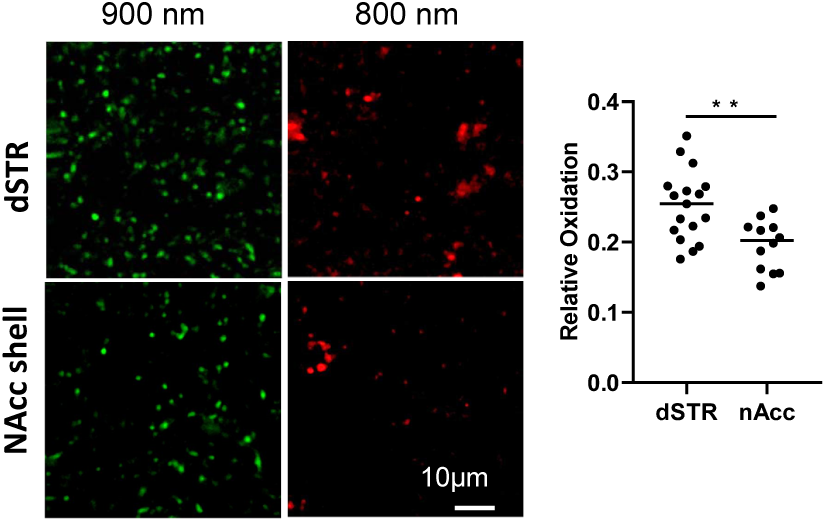
Oxidation level is higher in the dSTR than in the NAcc shell. **A**, Representative images roEGFP signals in the dSTR and NAcc excited at 900 nm and 800 nm. **B**, Plot of average oxidation level (800/900) in the dSTR (18 slices) and the NAcc (11 slices) from four 2-4-month-old mice, dSTR *vs* NAcc, *p* < *0.001*, Student t-test.

## Discussion

A large body of evidence from PD research has shown that mitochondrial dysfunction and oxidative stress is associated with PD (Guzman et al., 2010; Bose and Beal, 2016). Previous research has shown that basal OxPhos is higher in cultured SNc DA neurons compared to VTA DA neurons (Pacelli et al., 2015). Here, we report that evoked DA release in the dSTR is more dependent on OxPhos while evoked DA release in the NAcc shell is relatively more dependent on glycolysis. Consistent with the previous studies (Guzman et al., 2010; Pacelli et al., 2015), our results demonstrate an overall tendency for DA cell bodies and neurites associated with PD to be heavily reliant on mitochondria OxPhos in general while showing for the first time a relative glycolytic preference for DA release in the NAcc shell.

### Dorsal striatum prefers oxidative phosphorylation energy production for evoked DA release

Evoked DA release in the dSTR shows a clear preference for OxPhos for ATP production, while evoked DA release in the nAcc shell shows a higher preference for glycolytic ATP production compared to dSTR. The glycolytic preference for ATP production in the nAcc shell was apparent both when glycolysis was blocked (Fig. 4C) or when OxPhos was inhibited (Fig. 6C). Evoked DA release in the nAcc shell was maintained by OxPhos at around 50% when compared to control conditions (Fig. 5B, Fig. 6B, Fig. 9B). Under these conditions, the NAcc shell did not show a clear preference for either glycolysis or OxPhos for ATP production. Evoked release in the dSTR, however, clearly showed a preference for oxidative ATP production when OxPhos was selectively activated (Fig. 5A, Fig. 9A) and when OxPhos was inhibited (Fig. 6A).

Under glucose-deprived conditions, although the overall evoked DA release in the dSTR and the NAcc shell diminished to a similar degree over 40 min, the DA release in the NAcc shell was not diminished for the first ∼8 min compared to that in the dSTR (Fig. 3C). This trend was abolished when glycolysis inhibitor was administered (Fig. 4E), suggesting this is probably due to residual glycolysis.

### Monocarboxylate transport in the Striatum

Lactate is one types of monocarboxylate which must be exported out of cells as high levels of intracellular accumulation of lactate result in inhibition of glycolysis. However, some tissues, such as heart and brain, utilize lactate for cellular respiration and hence serve as an attractive candidate for alternative metabolic substrates during glycolytic inhibition.

Under normal condition when glucose is present, evoked DA release appears to be partially fueled by MCT in the dSTR but not the NAcc shell (Fig. 7, Fig. 9). This conclusion is further supported by the results of experiments with lactate in glucose-deprived condition. Just as with pyruvate in glucose-deprived conditions (Fig. 5C), evoked DA release in the dSTR was significantly greater than evoked DA release in the NAcc shell (Fig. 9C). Moreover, the trends of evoked DA release seen with lactate in glucose-deprived conditions are complementary with pyruvate in glucose-deprived conditions as well (Fig 5, Fig. 9). This demonstrates that the NAcc shell is not as effective and efficient as dSTR at using lactate or pyruvate for fuel source.

Under glucose-deprived conditions, it seems that both the dSTR and the NAcc shell utilize MCT as fuel to a similar extent (Fig. 9C). Recent evidence has shown that lactate shuttling involves astrocytes especially for axonal metabolism when energy demands are high, or glucose levels are low in the brain (Schurr et al., 1988; Magistretti and Allaman, 2018). Applying 4CIN in a glucose-deprived environment revealed a similar dependence on MCT to fuel evoked DA release in both the dSTR and the NAcc shell, consistent with the previous evidence. This evidence in conjunction with Fig. 8 and Fig. 4 also suggests that evoked DA release is more energetically demanding in the dSTR than the NAcc shell when glucose is present.

### Implications and Further Direction

The preference for oxidative energy production in the dSTR may be due to the overall difference in DA terminal density and complexity between the dSTR and the NAcc shell (Matsuda et al., 2009; Pacelli et al., 2015). The higher release of DA in the dSTR is due to a higher DA terminal density and a greater terminal complexity in the dSTR compared to the NAcc shell. Due to these factors, DA release in the dSTR is more energetically demanding than in the NAcc shell. To meet this increase in energy demand, energy production efficiency must be increased. Although both glycolysis and OxPhos produce ATP, ATP production via OxPhos is much more efficient than ATP production via glycolysis; increasing oxidative ATP production can significantly increase overall energy production without significantly increasing glucose consumption. Therefore increasing OxPhos relative to glycolysis supports DA release in the dSTR without increasing glucose consumption. A heavy reliance on OxPhos also allows the dSTR to be more flexible in its sources of fuel since OxPhos can be also fueled by lactate or ketones. This flexibility would allow DA release in the dSTR to be fueled even if energy demands increase or fasting/starvation occurs. According to the results presented here, the MCT system can fuel evoked DA release in the dSTR at similar levels to glucose, allowing DA release in the dSTR to be higher and more stable than in the NAcc shell given the same amounts of fuel.

The high dependence of dSTR on oxidative phosphorylation could in part help to explain the specific vulnerability of the DA terminals in the dSTR to degeneration in PD compared to the DA terminals in the NAcc shell. The dSTR utilizes OxPhos preferentially and the byproducts of OxPhos are reactive oxygen species that can lead to oxidative stress (Surmeier et al., 2011; Bose and Beal, 2016). Because the NAcc shell prefers glycolytic ATP production relavtively, lower levels of oxidative stress and neurodegeneration would occur. But the protection may go further than that. The NAcc shell utilizes a relatively higher level of glycolysis to fuel the evoked DA release. Lactate, a byproduct of glycolysis, can activate pathways associated with cellular survival, homeostasis, angiogenesis, and overall neuroprotection (Yang et al., 2014; Brooks, 2018; Magistretti and Allaman, 2018). Upregulation of glycolysis is neuroprotective in a Drosophila model of amyotrophic lateral sclerosis (Manzo et al., 2019). Likewise, recent research in clinical models has shown that enhancing glycolysis attenuates PD symptoms (Cai et al., 2019). Further study to examine the different expression levels of the metabolic genes and proteins may uncover the underlying mechanisms driving the different preferences of ATP-producing pathways in the dSTR and the NAcc shell.

## Acknowledgments/Conflict of interest disclosure

We thank Dr. D. James Surmeirer for providing the TH-mito-roGFP mice. This work was supported by the National Institute of Neurological Disorders and Stroke (NINDS) (NS098393 and NS097530 to H.Z.) and the Department of Neuroscience at Thomas Jefferson University (Startup Funds to H.Z.). The authors declare no competing financial interests.

